# Transcutaneous electrical nerve stimulation modulates corticospinal excitability while preserving motor unit discharge properties during isometric contractions

**DOI:** 10.1101/2022.03.10.483702

**Authors:** Julio Cesar Hernandez-Pavon, Simon Avrillon, Grace Hoo, Jose L. Pons

## Abstract

**Aim:** Transcutaneous electrical nerve stimulation (TENS) aims to supplement sensory feedback to improve force steadiness or motor function. In this study, we directly assessed potential changes in corticospinal excitability and motor unit discharge characteristics from the first dorsal interosseous (FDI) muscle due to TENS by using transcranial magnetic stimulation (TMS) and high-density surface electromyography (HDsEMG).

**Methods:** Eleven healthy young adults performed a series of submaximal isometric index abductions. We estimated i) motor evoked potential (MEP) amplitudes, ii) persistent inward current amplitudes (PIC, i.e., delta F), iii) motor unit recruitment thresholds and discharge rates, and iv) common synaptic input to motor units before and after TENS.

**Results:** TENS did not affect force steadiness (2.5 ± 0.9% and 3.3 ± 1.9% (*p* = 0.010)). MEP amplitudes decreased at 110% of the resting motor threshold (rMT; 0.72 ± 0.66 mV *vs*. 0.59 ± 0.63 mV; *p* < 0.001), increased at 130% rMT (1.18 ± 1.10 mV *vs*. 1.41 ± 1.29 mV; *p* < 0.001). Delta F increased after TENS (3.7 ± 2.2 pps *vs*. 4.5 ± 2.6 pps; *p* = 0.010). We did not find a change in the level of common synaptic input or in the temporal variability of motor unit discharge rates after the session of TENS.

**Conclusion:** These results suggest that TENS can modulate corticospinal excitability through supraspinal and spinal processes and, thus act as a priming technique. At the same time, TENS does not generate short-term changes in the neural control of force in young, healthy adults.

## 1. INTRODUCTION

The central nervous system controls muscle force by recruiting motor units and modulating their discharge rates. Most motor units from the same pool display common modulation of their discharge rate during isometric tasks due to the high proportion of common synaptic input they receive [1, 2]. Therefore, the slight variations of force during submaximal isometric contractions are highly correlated to the common variations in motor unit discharge rates [1, 3]. Several studies have documented how aging and neurological diseases, such as essential tremor or multiple sclerosis, increase the temporal variability of the common synaptic input [4–7]. This leads to decreased force steadiness, reduced manual dexterity, or decreased walking performance [5–8].

One way to compensate for impaired motor unit activation is to supplement the motor and sensory neural commands with transcutaneous electrical nerve stimulation (TENS) [9]. TENS consists of applying an electrical current at low intensity and high frequency to target the recruitment of sensory fibers, which have a lower rheobase than motor fibers [10, 11]. For example, Hamilton et al. [12] found that TENS applied over the biceps brachii increased the electromyography amplitude recorded over the contralateral biceps brachii. Alternatively, the stimulation of the cutaneous afferent fibers of the ankle for 20 minutes increased the spinal excitability of motor neurons innervating the soleus muscles [13]. Besides the acute changes at the spinal level, TENS can also reach supraspinal centers [14, 15]. For example, the chronic stimulation of hand afferent fibers led to an increase in corticospinal excitability assessed with transcranial magnetic stimulation (TMS) [16]. Importantly, how TENS and its effect on spinal and supraspinal excitability impact the neural control of force during isometric contractions is unknown.

It is now possible to identify numerous motor units from electromyographic signals recorded with a high density of electrodes [17]. This method allows researchers to simultaneously study the dynamics of many motor units and thus describe the common synaptic input to motor units and its link with force steadiness [1, 2, 7]. Taking advantage of the unique representation of motor unit action potentials through the multiple electrodes, we can also track the same motor units across time and conditions [18, 19], which offers a formidable tool to assess the potential effect of TENS on motor unit discharge properties [12] and to determine how TENS impacts motor unit excitability during isometric contractions [20]. Indeed, the transformation of synaptic input into motor unit discharge trains is mediated by neuromodulation processes through the activation of persistent inward currents (PIC) [21, 22]. PIC can amplify and prolong synaptic input to motor units during steady contractions [21]. Importantly, the stimulation of sensory pathways with electrical stimulation or tendon vibration acutely triggers these PIC [10, 23]. Moreover, experimental and simulation studies have found that sensory pathways generating inhibition dramatically altered the amplitude of PIC during isometric contractions [24, 25]. In this way, a session of repeated stimulation of sensory pathways with TENS may also impact the amplitude of PIC, which remains unknown.

This study investigated how a single session of TENS in healthy participants impacts corticospinal excitability and motor unit discharge properties from the first dorsal interosseous (FDI) during isometric index abductions. The TENS aimed to stimulate cutaneous and Ia afferent fibers from the hand using a current intensity below the motor threshold delivered at 100 Hz. We assessed the change in corticospinal excitability by measuring the amplitude of TMS motor evoked potentials (MEPs). Then, we identified and tracked motor unit discharge times during isometric contractions by decomposing high-density surface electromyographic (HDsEMG) signals with a blind source separation algorithm [18, 26]. We estimated the change in PIC amplitude using the delta F method. We also evaluated the impact of TENS on the neural control of force by comparing motor unit recruitment thresholds, discharge rates, the level of common input to motor units, and the temporal variability of the smoothed discharge rate of the cumulative spike trains before and immediately after the TENS session. We hypothesized that TENS would increase corticospinal excitability with increases in MEP and PIC amplitudes. We also hypothesized that TENS would not significantly impact the neural control of force and force steadiness in our population of young, healthy adults, as reported in a previous work [27]. Thus, TENS would act as a priming technique to increase corticospinal excitability, which may be useful for clinical applications.

## 2. MATERIALS AND METHODS

### 2.1 Participants

Eleven healthy young volunteers (mean ± standard deviation; age: 31 ± 10 years, height: 175 ± 10 cm, body mass: 76 ± 15 kg; 5 women, all right-handed) participated in the study. All experimental procedures were approved by the Northwestern University IRB (IRB#: STU00211930; Clinical Trials: NCT04501133) and were in accord with the Declaration of Helsinki. All participants gave their written informed consent before the experiments, and none of them had neither brain lesions/neurological disorders nor contraindication to electrical stimulation or TMS [28].

### 2.2 Experimental design

Participants sat in a chair with adjustable height and hip angle to optimize their comfort. A force sensor (MLP-200, Transducer Techniques, CA, USA) was fixed and aligned with the right armrest. The right forearm and arm were strapped to minimize any movements during the contractions.

The experimental tasks consisted of a series of isometric index finger abduction against the center of the force sensor with or without electrical stimulation. The force signal was digitized at 2048 Hz using the HDsEMG acquisition system (Quattrocento, OT Bioelettronica, Turin, Italy). The level of force produced by the participant was displayed on a screen positioned in front of the participant. We recorded surface electromyographic signals of the first dorsal interosseous (FDI) muscle during these tasks with a grid of 64 electrodes (GR04MM1305, Quattrocento, OT Bioelettronica, Turin, Italy).

We divided the session into three blocks with 90 s of rest in between. The first and the third blocks consisted of pre and post measurement of i) corticospinal excitability, ii) persistent inward currents (PIC), iii) motor unit recruitment thresholds and discharge rates, and iv) common input to motor units (**Figure 1**). The second block consisted of a series of nine submaximal isometric contractions during which we delivered transcutaneous electrical nerve stimulations (TENS) (**Figure 1**). We ultimately aimed to determine whether the TENS session elicits changes in the neural control of force at the spinal and supraspinal levels by comparing measurements from the first and third blocks.

**FIGURE 1.**
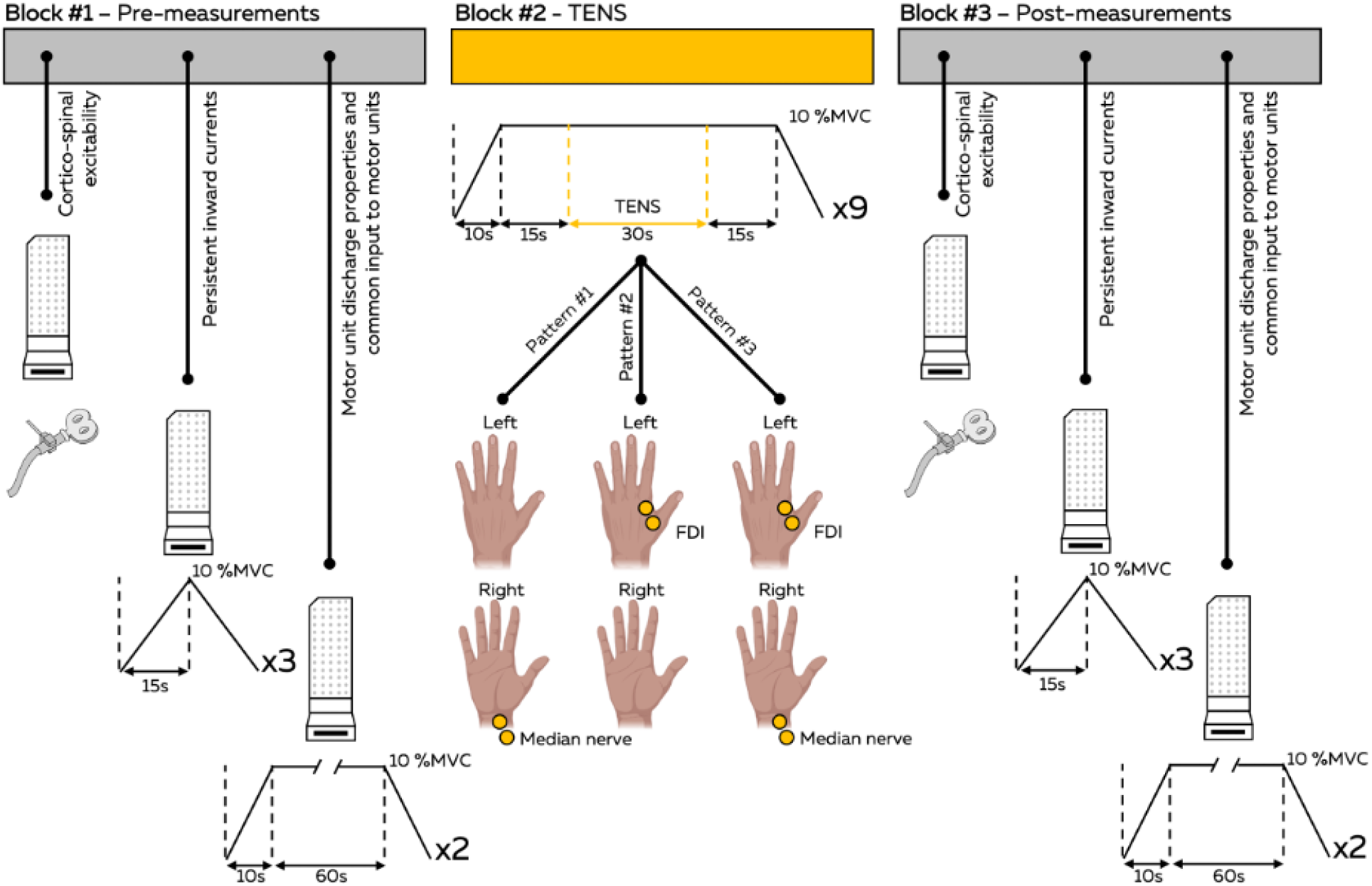
Experimental timeline for a participant. Each participant performed three blocks with 90 s of rest in between, consisting of two blocks of pre-and post-measurements (Blocks #1 and #3) separated by a transcutaneous electrical nerve stimulation (TENS) session. We assessed the corticospinal excitability by pairing transcranial magnetic stimulation (TMS) with high-density surface electromyography (HDsEMG) recordings using a grid of 64 electrodes. We estimated persistent inward currents by identifying motor unit discharge times from HDsEMG signals during a 30 s triangular isometric contraction up to 10% of the maximal voluntary contraction (MVC). We assessed changes in motor unit discharge properties and common input to motor units by identifying motor unit discharge times from HDsEMG signals during an 80 s trapezoidal isometric contraction with 10 s ramp-up and -down and a 60 s plateau set at 10% of the MVC. During the TENS session, we activated sensory afferent fibers to increase proprioceptive feedback and change the excitability of the central nervous system. We randomized three complementary stimulation patterns: Pattern #1, stimulation of the right median nerve; Pattern #2, stimulation of the left first dorsal interosseous (FDI) muscle belly; and Pattern #3, stimulation of both the median nerve and the left FDI muscle belly. Each stimulation pattern was delivered during a 30 s window centered within the middle of the plateau.

To assess changes in corticospinal excitability, participants positioned their right hand with the palm inward and the forearm in a neutral position. 20 TMS biphasic pulses were administered to the left cortical representation of the FDI muscle per intensity at 110%, 120%, and 130% of the resting motor threshold (rMT). The MEPs were evoked at an inter-stimulus interval of 5 s, and the order of the TMS intensities was randomized. Then, after a standardized warm-up, the participant performed three maximal isometric voluntary contractions (MVC) for 10 s with 90 s of rest in between. We applied a moving-averaged window of 250 ms over the force signals and considered the maximal value obtained across the three MVC as the maximal force. To assess changes in motor unit recruitment thresholds and discharge rates and in common input to motor units, the participants performed two submaximal isometric abductions of the index finger during 80 s with 30 s of rest in between. During this task, participants matched a trapezoidal target with 10 s ramp-up and ramp-down and a 60 s plateau set at 10% of the MVC. To assess changes in persistent inward currents, the participants performed three submaximal isometric abductions of the index finger during 30 s with 30 s of rest in between. During this task, participants tracked a triangular target with 15 s ramp-up and ramp-down with a maximum set at 10% of the MVC (rate of 0.66% MVC.s^-1^).

During the block of TENS, participants performed nine 80 s submaximal isometric abductions of the index finger with 10 s ramp-up and ramp-down, and a 60 s plateau at 10% of the MVC separated by 30 s of rest. TENS was applied during a window of 30 s centered over the middle of each plateau. A total of 19 contractions at 10% of the MVC was performed per participant during the entire session.

### 2.3 Transcutaneous electrical nerve stimulations (TENS)

The transcutaneous electrical nerve stimulation (TENS) session aimed to activate sensory fibers during repeated submaximal isometric contractions to provide additional proprioceptive feedback and change the excitability of the central nervous system. It is noteworthy that a current intensity below the motor threshold can specifically activate sensory afferent fibers, as their rheobase is lower than the motor fibers [10]. Also, the position of the stimulation over the nerve or the muscle belly can favor the recruitment of Ia afferent fibers or cutaneous afferent fibers, respectively [12, 29, 30]. In this study, we stimulated the median nerve on the right hand and the FDI muscle belly on the left hand. We first used handheld dry electrodes with an anode and a cathode of 12 mm diameter separated by 50 mm and connected to a bipolar constant current stimulator (Digitimer DS5, Digitimer, UK) to find the optimal location that selectively activated the thenar eminence on the right side and the FDI muscle on the left hand. Then, the skin was cleaned with alcohol, and two round electrodes were placed over the skin (PALS electrodes with a diameter of 2.5 cm, Axelgaard, USA). We then found the sensory threshold, i.e., the minimal current where the participant could feel the stimulation, and the motor threshold, i.e., the minimal current intensity where the muscle contraction was visible. The level of intensity was set to the highest current intensity below the motor threshold. When this current intensity was too painful for the participant, we lowered the current intensity until reaching a tolerable intensity, with a minimal intensity set to 1.5 × the sensory threshold. During the session, the stimulation was delivered at a frequency of 100 Hz with a pulse duration of 400 μs. We randomized three complementary stimulation patterns: i) stimulation of the right median nerve, ii) stimulation of the left FDI muscle belly, iii) stimulation of both the median nerve and the left FDI muscle belly. Each stimulation pattern was delivered during three blocks of 30 s, for a total of nine blocks of 30 s during the entire session (**Figure 1**). Previous work showed that the activation of the median nerve, mostly composed of sensory fibers [31], impacted the activation of the FDI muscle [29]. Similarly, the TENS of a muscle increased motor unit output of the same contralateral muscle [12]. We were, therefore, confident that all the TENS patterns would similarly impact the motor unit output of the targeted FDI muscle.

### 2.4 High-density surface electromyography (HDsEMG) recordings

We recorded HDsEMG signals from the FDI muscle using a two-dimensional adhesive grid of 64 electrodes (13 × 5 electrodes with one electrode absent from a corner, inter-electrode distance: 4 mm, OT Bioelettronica, Turin, Italy). The grid was aligned with the direction of the muscle from the first metacarpal bone to the proximal phalanx of the index finger. Before electrode application, the skin was shaved and cleaned with an abrasive pad and alcohol. The grid was placed on the skin using adhesive foam layers with cavities filled with conductive paste over each electrode to optimize skin-electrode contact. Strap electrodes dampened with water were placed around the wrist of the tested arm as ground and reference electrodes. The HDsEMG signals were recorded in monopolar mode, bandpass filtered (10–900 Hz), and digitized at a sampling rate of 2048 Hz using a multichannel acquisition system (EMG-Quattrocento; 400-channel EMG amplifier, OT Bioelettronica, Turin, Italy).

### 2.5 TMS and Neuronavigation

Transcranial magnetic stimulation (TMS) was applied with a figure-of-eight coil (C-B60, MagVenture, Farum, Denmark) connected to a TMS stimulator (MagPro X100 with MagOption, MagVenture, Farum, Denmark). TMS was targeted with a neuronavigation system (Localite, St Augustin, Germany) using T1-weighted MRIs (T1-weighted MRI, MEMPRAGE at 1 mm isotropic resolution) previously recorded in a 3T scanner (Siemens Prisma 3T, Siemens Medical Solutions, Erlangen, Germany) for each participant. This allowed us to easily identify the cortical representations of the FDI in the left motor cortex, i.e., the brain location where contractions of the right FDI are encoded, and to keep the TMS coil at the exact same location and orientation during the session. The primary motor cortex representations of the FDI were identified by moving the TMS coil over the central sulcus/precentral gyrus while keeping the coil orientation perpendicular to the local curvature of the central sulcus. Once the optimal location of the coil was found, we determined the rMT as the lowest TMS intensity evoking MEPs with an amplitude >50 μV peak-to-peak in at least five out of ten consecutive trials. Participants wore earplugs to protect their hearing from the TMS coil clicks during the stimulation. At each TMS pulse, a single logic pulse was sent to the HDsEMG amplifier to synchronize HDsEMG signals with the TMS.

### 2.6 Data processing

#### 2.6.1 Motor evoked potentials (MEPs)

The monopolar EMG signals from the 64 channels were bandpass filtered between 10-500 Hz with a third-order Butterworth filter. The 64 HDsEMG channels were segmented over 100 ms after each stimulation and averaged across the 20 trials per intensity. The 64 averaged MEPs were then displayed for visual inspection. Channels with artifacts or low signal-to-noise ratio were discarded from further analysis. The peak-to-peak amplitudes were calculated within a window from 26 to 40 ms after the TMS trigger.

We applied two approaches to assess the changes in MEP amplitude before and after the TENS session. In the first approach, we compared the MEP values from the entire grid. To this end, we considered the channels which were not discarded in all the intensities before and after the TENS session (**Figure 2A**). In the second approach, we defined a region of interest that only kept the channels over the muscle belly, where the MEP amplitudes should be higher [32]. To this end, we considered the map of MEP amplitudes as a topographical map where *i* and *j* are the coordinates of the electrodes and *z* is the peak-to-peak amplitude of the MEP value. Then, we applied an h-dome transform that considered all the peaks above a height *h* and refined the results with a morphological opening to avoid the segmentation of fractioned and small areas, as described in [32]. We considered the electrodes in the region of interest (ROI) in all the intensities before and after the TENS session (**Figure 2A**).

**FIGURE 2.**
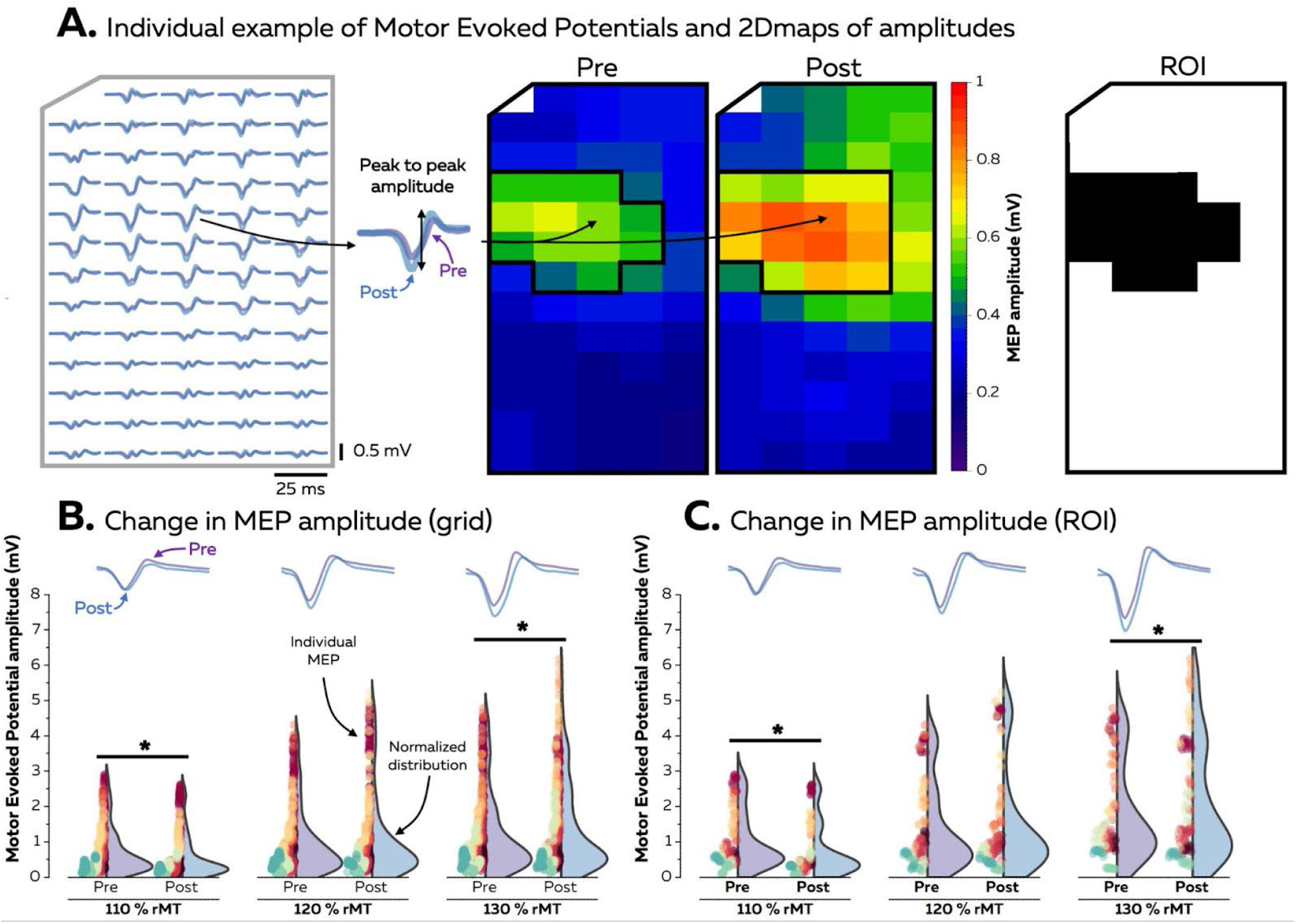
Analysis of MEPs evoked with TMS. **A**. Individual example of averaged MEP recorded at 110% rMT within a grid of 64 electrodes. The purple traces represent the MEPs recorded before the TENS session, and the blue traces represent the MEPs recorded after the session. The peak-to-peak amplitudes were assessed from 26 to 40 ms after the TMS for all the channels. Two complementary analyses were performed to compare MEP amplitudes before and after the TENS session. We first compared the MEP amplitudes from the channels that were not discarded during all the conditions for each participant **(B)**. Then, we used the two-dimensional images of MEP amplitudes within the grid to define a region of interest (ROI) over the muscle belly, i.e., where the MEP amplitudes are the highest. We compared the MEP amplitudes from the channels that were in the ROIs during all the conditions for each participant **(C)**. On panels **B** and **C**, the purple traces represent the averaged MEPs recorded before the session, and the blue traces represent the averaged MEPs recorded after the session. All the MEP amplitudes are displayed with a different color per participant. The distributions are normalized by the maximal count of MEP amplitudes. * Denotes a significant difference between pre and post values for a given intensity.

#### 2.6.2 HDsEMG signals decomposition

The monopolar EMG signals from the 64 channels were bandpass filtered between 20-500 Hz with a second-order Butterworth filter. After visual inspection, channels with a low signal-to-noise ratio or artifacts were discarded. The HDsEMG signals were then decomposed into motor unit spike trains using convolutive blind source separation as previously described [26]. The EMG signals were first extended and spatially whitened. Then, we applied a fixed-point algorithm that maximized the sparsity to identify the sources of the EMG signals, i.e., the motor unit spike trains. The spikes were separated from the noise using K-mean classification and a second algorithm refined the estimation of the discharge times by minimizing the coefficient of variation of the inter-spike intervals. After 150 iterations, duplicates, considered as motor unit spike trains that share more than 30% of their discharge times, were discarded from the analysis. Finally, all the spike trains were visually checked for false positives and false negatives. We only kept the motor units that exhibited a pulse-to-noise ratio > 30 dB for further analysis, thus ensuring a sensitivity higher than 90% and a false-alarm rate lower than 2% [33].

#### 2.6.3 Motor unit tracking

We performed separate HDsEMG signal decomposition for each condition, leading to identifying different pools of motor units before and after the TENS session. Therefore, we tracked the motor units over time to compare recruitment threshold and discharge rate, common input to motor units, and persistent inward current on pools with similar properties. To this end, we matched motor units using the distribution of action potential shapes within the entire grid (**Figure 4A**) [18, 19]. Specifically, the motor unit action potential shapes were identified using the spike-triggered averaging technique. We used the discharge times as a trigger to segment and average the HDsEMG signal over a window of 50 ms for each channel. As the position of the action potentials within the window may differ between motor units, we used the local maxima, i.e., the action potential with the highest amplitude, as the center of the window. Finally, we concatenated the motor unit action potentials from the 64 channels, and we performed a 2D cross-correlation between motor units. Pairs with a correlation coefficient R higher than 0.8 were considered as matches [18, 19]. The matches were visually checked by an experienced operator (SA) to guarantee the similarity of the motor unit action potential shapes. We compared the recruitment thresholds, i.e., the normalized joint torque at the time the motor unit begins to discharge, and the discharge rates of these motor units before and after the session.

**FIGURE 3.**
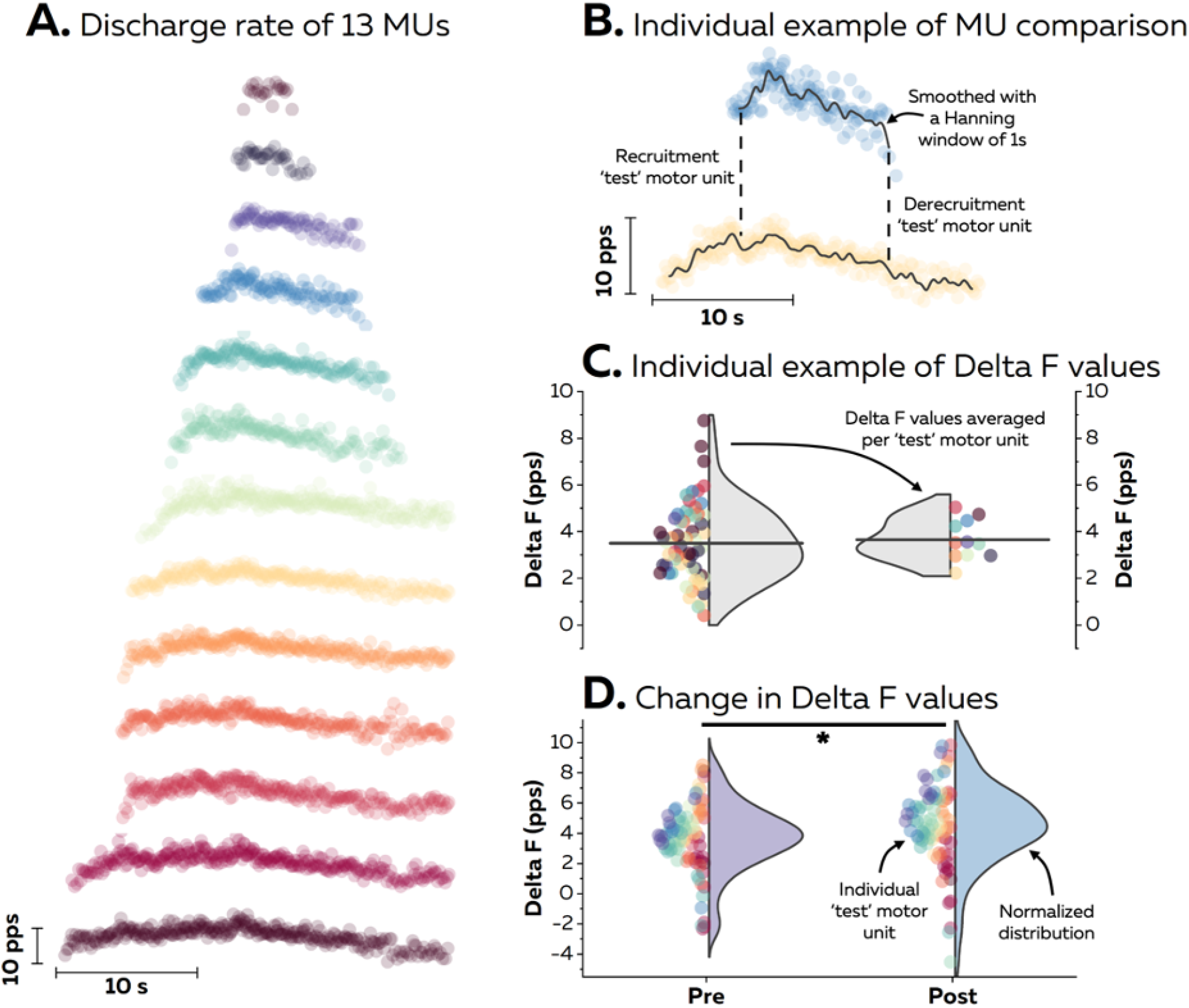
Analysis of persistent inward currents. **A**. Individual example of instantaneous motor unit discharge rates during a triangular contraction with a peak set at 10% of the MVC. Motor units were paired to estimate the delta F when their recruitment and derecruitment times were separated by 1 s and 1.5 s, respectively. **B**. For each pair, the delta F value was calculated as the difference in smoothed instantaneous discharge rate of the ‘control’ motor unit at the recruitment and derecruitment times of the ‘test’ motor unit. **C**. Delta F values were averaged per ‘test’ unit. The color code on panels **A** and **C** represents the same motor units. **D**. Delta F values for the entire population of motor units before and after the session. All the motor units are displayed with a different color per participant. The distribution is normalized by the maximal count of motor units.

**FIGURE 4.**
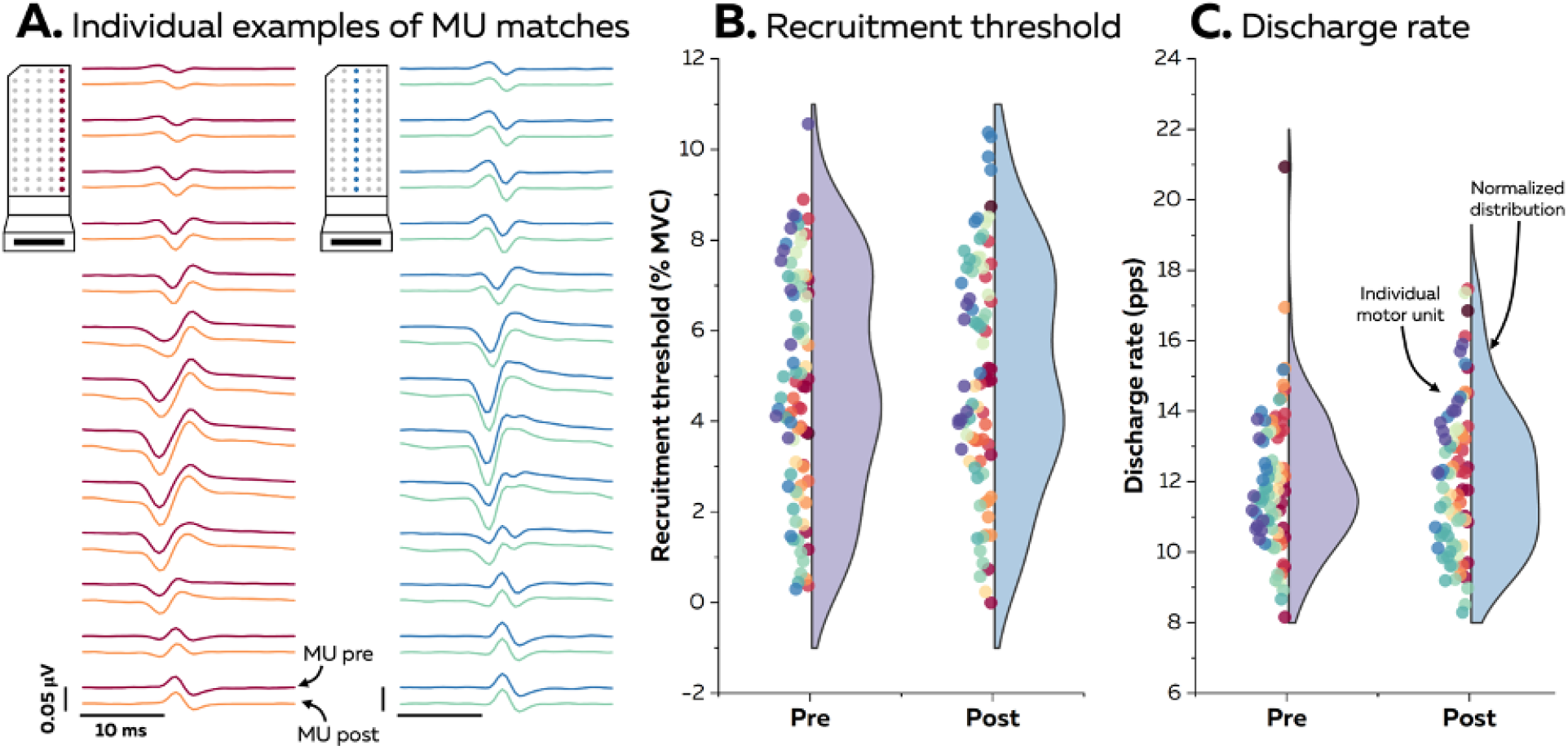
Analysis of motor unit discharge properties. **A**. Example of motor unit tracking for two individual motor units. Motor unit action potential shapes are displayed along a column of 13 electrodes. A two-dimensional cross-correlation was computed on the motor unit action potentials concatenated over the entire grid to track motor units. Motor units were matched when the correlation coefficient exceeded R = 0.8. Recruitment thresholds **(B)** and discharge rates **(C)** are displayed for individual motor units. All the motor units are displayed with a different color per participant. The distribution is normalized by the maximal count of motor units.

#### 2.6.4 Estimation of Persistent Inward Currents (PIC)

The relation between motor unit input and output is nonlinear due to neuromodulation processes with the activation of PIC. PIC can amplify and prolong excitatory synaptic input to motor neurons [21]. The effect of PIC on motor units discharge rates is estimated using the hysteresis between motor unit discharge rates at recruitment and derecruitment of a ‘test’ motor unit (**Figure 3B**) [34]. This method, called delta F, consists of calculating the difference in discharge rate of the ‘control’ motor unit at the recruitment and derecruitment times of the ‘test’ motor unit (**Figure 3B**). The ‘test’ motor unit has higher recruitment and derecruitment thresholds than the ‘control’ motor unit. Here, we used a threshold of 1 s for the time difference in the recruitment threshold and 1.5 s for the time difference in the derecruitment threshold [35]. The instantaneous discharge rates of all the motor units were smoothed using a Hanning window of 1 s [35]. Delta F values were averaged per ‘test’ motor unit [36].

#### 2.6.5 Motor unit coherence analysis

The force steadiness during isometric contractions depends on the oscillations in discharge rate in the bandwidth of force fluctuation (0-5 Hz) shared by many motor units from the same pool [7]. The strength of this common input to motor units is classically inferred from coherence analysis. This analysis represents the correlation between two motor unit smoothed discharge rates at given frequencies, with 0 indicating no correlation and 1 indicating a perfect correlation. Here, we calculated the pooled coherence on two equally sized groups of cumulative spike trains (**Figure 5A**). Matched motor units that discharged continuously, i.e., without pauses > 250 ms, were used to compute the coherence over a window of 30s before and after the TENS session. The number of motor units in each of the two groups varied from one to the maximum number (half of the total number of identified units). For each iteration, the motor units used to compute the cumulative spike trains were permuted with up to 100 random permutations per iteration. We calculated the coherence using Welch’s periodogram with nonoverlapping Hanning windows of 1 s. We compared the area under the coherence curve between groups of two motor units (the minimum number of identified motor units being four) from 0 to 5 Hz as an indicator of the strength of common input before and after the TENS session. We also compared the rate of increase between groups of one and groups of two motor units and considered this increase as an estimate of the proportion of common input [37].

**FIGURE 5.**
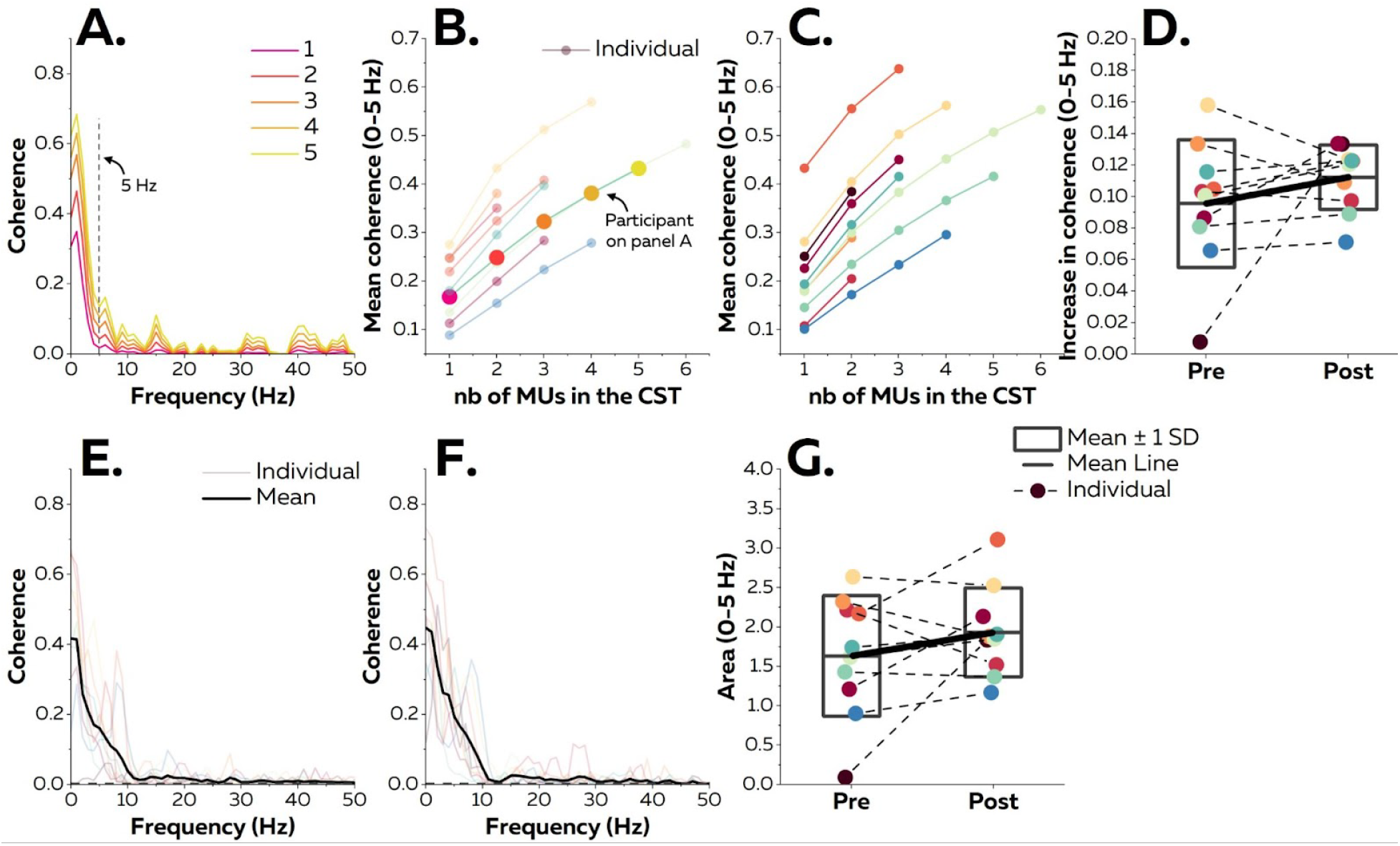
Analysis of common input to motor units. **A**. We performed a coherence analysis on two equally sized groups of cumulative spike trains (CST). In this example, the coherence analysis was performed on groups of one (pink trace) to five motor units (yellow trace). As the control of force depends on the common slow oscillations of discharge rates in the bandwidth of force fluctuation, we focused our analysis on the 0-5 Hz bandwidth (Panels **B, C, D, G**). The mean coherence between 0 and 5 Hz increases as the number of motor units used to calculate the cumulative spike train (CST) increases before (**B**) and after (**C**) the session. **D**. Rate of increase in mean coherence from 0 to 5 Hz before and after the session. Each scatter represents one participant. The boxes represent the mean ± standard deviation. **E** and **F** display the level of coherence estimated between groups of two motor units. We estimated the strength of common input using the area under the coherence curve from panels **E** and **F (G)**.

### 2.7 Statistical analysis

To inspect the normal distribution of each dataset, we plotted normal probability plots and histograms. Data that deviated from the normal distribution were transformed to display a normal distribution. Values are reported as means ± standard deviation.

All statistical analyses were implemented in RStudio. It is noteworthy that we considered all the data points for each analysis instead of averaging the values per participant. For example, we compared the recruitment thresholds of 79 motor units instead of nine averaged recruitment thresholds to better represent the effect of the session on the total population of motor units. To this end, we used linear mixed effect models (LMM) implemented in the R package *lmerTest* with the Kenward-Roger method to estimate the denominator degrees of freedom and the *p*-values.

Importantly, this method takes account of the dependence of data points within each participant due to, for example, common input to motor units from the same pool [38].

To compare MEP amplitudes from the entire grid and the region of interest before and after the session, we used a LMM with intensity (110%, 120%, and 130% rMT) and time (pre, post) as fixed effects and participants as a random effect. To compare recruitment thresholds, discharge rates, areas under the coherence curve, rate of increase in coherence, and delta F values before and after the TENS session, we used a LMM with time (pre, post) as a fixed effect and participants as a random effect. When necessary, we performed multiple comparisons using the R package *emmeans*, which adjusts the *p*-value using the Tukey method. The significant level was set at 5% (*p* < 0.05).

## 3. RESULTS

Participants successfully tracked the force target at 10% MVC with normalized values of 9.7 ± 1.3% MVC during the plateau of 60 s isometric contractions. Force steadiness, i.e., the coefficient of variation of force during the plateau, did not significantly change after the TENS session, with respectively 2.5 ± 0.9% and 3.3 ± 1.9% (*p* = 0.010). We then compared i) MEP amplitudes, ii) PIC amplitudes, iii) motor unit recruitment thresholds and discharge rates, and iv) common input to motor units before and immediately after the session of TENS.

### 3.1 Change in MEP amplitude

Data from participants #3 and #4 were discarded from the TMS analysis as we were not able to evoke MEPs in the FDI. We combined TMS and HDsEMG to record MEP amplitudes over the 64 channels of the grid at 110% of resting motor threshold (rMT), 120% rMT, and 130% rMT (**Figure 2A**). When considering MEP amplitudes over the entire grid, we found a significant Intensity x Time interaction (**Figure 2B**; F = 42.869; *p* < 0.001). Specifically, the MEP amplitude decreased at 110% rMT (pre: 0.72 ± 0.66 mV *vs*. post: 0.59 ± 0.63 mV; *p* < 0.001) and increased at 130% rMT (pre: 1.18 ± 1.10 mV *vs.* post: 1.41 ± 1.29 mV; *p* < 0.001). No differences were found between pre and post MEP amplitudes at 120% rMT (*p* = 0.058). We then only considered a region of interest over the muscle belly using an image-based segmentation (**Figure 2A**; see the Material and Methods section for further details). We found similar results with a significant Intensity x Time interaction (**Figure 2C**; F = 22.221; *p* < 0.001). The MEP amplitudes decreased at 110 % rMT (pre: 1.00 ± 0.79 mV *vs.* post: 0.75 ± 0.75 mV; *p* < 0.001), increased at 130% rMT (pre: 1.76 ± 1.35 mV *vs.* post: 2.08 ± 1.63 mV;*p* = 0.009) while we did not find any significant difference at 120% rMT (*p* = 0.96).

### 3.2 Change in Persistent Inward Currents (PIC)

We used an algorithm based on convolutive blind source separation to decompose HDsEMG signals into motor unit spike trains (**Figure 3A**). Note that we matched motor units before and after the session of TENS to compare groups of motor units with similar properties (see the Material and Methods section for further details). We identified a total of 267 motor units during the triangular isometric contractions before and after the TENS session, with an average of 12 ± 4 motor units per contraction. We tracked 103 motor units (9 ± 4 motor units per participant) with a cross-correlation coefficient of 0.93 ± 0.05. Then, we estimated the impact of PIC on motor unit discharge rates using the delta F method, which consists of comparing the discharge rates of a ‘control’ – low threshold – motor unit at the recruitment and derecruitment times of a ‘test’ – high threshold – motor unit (**Figure 3B**; [34]). We averaged all the delta F values per ‘test’ motor unit (**Figure 3C**). A significant time effect was found on delta F values (F = 6.781; *p* = 0.010). Specifically, the delta F values increased from 3.7 ± 2.2 pps to 4.5 ± 2.6 pps after the TENS session (**Figure 3D**).

### 3.3 Change in Motor Unit discharge properties

A total of 235 motor units were identified from HDsEMG signals during the isometric trapezoidal contractions at 10% of the MVC, with an average of 11 ± 4 motor units per contraction. 79 motor units were matched between pre and post TENS, with a cross-correlation coefficient of 0.93 ± 0.05 (**Figure 4A**). There was neither time effect for the motor unit recruitment thresholds (**Figure 4B**; F = 0.539; *p* = 0.46) nor the discharge rates (**Figure 4C**; F = 0.494; *p* = 0.48). Specifically, motor units began to discharge at 4.8 ± 2.5% MVC and 5.1 ± 2.6% MVC before and after the TENS session, respectively. Motor units discharged at a rate of 12.0 ± 2.0 pps and 12.1 ± 2.1 pps before and after the session, respectively.

### 3.4 Change in common input to motor units

73 motor units that continuously discharged during 30 s were matched before and after the TENS session, with an average of 7 ± 3 motor units per participant. We estimated the level of common input to motor units using a coherence analysis. As the control of force depends on the common slow oscillations of discharge rates in the bandwidth of force fluctuation, we focused our analysis on the 0-5 Hz bandwidth (**Figure 5A**). The coherence analysis was only performed on ten participants, as only one motor unit was matched for participant #1. There was no time effect on the area under the coherence curve between 0 and 5 Hz (F = 1.666; *p* = 0.22), with averaged values of 1.63 ± 0.76 and 1.93 ± 0.57 before and after the TENS session, respectively (**Figure 5G**). We then estimated the proportion of common input using the increase in mean coherence from 0 to 5 Hz between groups of one motor unit and groups of two motor units (**Figures 5B–5D**). There was no time effect on the rate of increase in coherence (F = 1.216; *p* = 0.30), with averaged values of 0.10 ± 0.04 and 0.11 ± 0.02 before and after the session, respectively. Finally, we estimated the temporal variability of the discharge rate of the cumulative spike train at low frequencies (i.e., 2.5 Hz). To this end, we smoothed the cumulative spike train by convoluting the binary signals (motor unit spike times) with a 400 ms Hanning window. The coefficient of variation of the smoothed discharge rates did not significantly change after the session (8.6 ± 3.3% and 9.6 ± 2.2% before and after the session, respectively; *p* = 0.42).

## 4. DISCUSSION

In this study, we used TMS and HDsEMG to investigate how a TENS session impacts corticospinal excitability and motor unit discharge properties during submaximal isometric abductions of the FDI muscle. We found that TENS modulated the excitability of motor neurons with changes in MEP amplitudes and delta F values. Conversely, motor unit discharge properties and the level of common input to motor units did not change. These results suggest that TENS can modulate corticospinal excitability without impacting the dynamics of the neural control of force during isometric contraction in healthy participants. In this way, TENS may act as a priming technique with potential applications in a clinical setting.

TENS aimed at activating cutaneous and proprioceptive afferent feedback to increase corticospinal excitability of the FDI muscle in healthy participants. Simulations and experimental recordings showed that TENS allows recruiting motor units acutely at the spinal level according to the size principle while minimizing muscle fatigue [10, 11, 39]. In rehabilitation settings, this method associated with motor training tended to improve hand dexterity in patients post-stroke [40] or with multiple sclerosis [41]. However, to date, no consensus exists regarding the optimal stimulation parameters, i.e., pulse width and frequency, that increase corticospinal excitability and improve motor function. Both wide pulses (1 ms [10]) and narrow pulses (200 μs [12]) increased motor unit outputs in the targeted muscle. In a clinical setting where patients with multiple sclerosis participated in TENS and training sessions to improve motor function, Almukass et al. [42] found similar clinical outcomes with stimulators delivering narrow (200 μs) and wide (1 ms) pulses. When considering the frequency, stimulations at 100 Hz were the most effective to increase motor unit output [10, 11] or increase corticospinal excitability [43]. In line with these results, we used TENS with a pulse duration of 400 μs delivered at 100 Hz to elicit changes in corticospinal excitability and motor unit output.

We first showed that TENS modulated corticospinal excitability with a decrease of MEP amplitudes at 110% rMT, no changes at 120%, and an increase at 130% rMT (**Figure 2**). The increase at 130% rMT is consistent with a previous study that found an increase of MEP amplitudes in the FDI muscle after sessions of TENS from 120% to 160% rMT, with no changes below 120% rMT [16]. Alternatively, Mima et al. [44] observed a short-term reduction of MEPs amplitude of the FDI evoked at 115% rMT. It is noteworthy that the differences in changes in MEP amplitudes between 110% rMT and 130% rMT are unlikely to be TENS time dependent as we randomized TMS intensities across participants. This difference may arise from the cortical brain area stimulated by the TMS, which increases with higher intensities of the stimulation [45, 46]. Previous work had shown that the area of the cortical representation of the FDI muscle decreased when an anesthetic block of the median nerve suppressed cutaneous feedback from the hand [29]. On the contrary, the area of cortical representations of a muscle can increase after the stimulation of peripheral nerves [47]. Therefore, the stimulation of a larger brain area surrounding a larger cortical representation of the FDI at 130% rMT may explain our results. Locally, TENS may induce a decrease in cortical excitability, as shown at 110% rMT. In this way, GABAergic mechanisms are likely to play a significant role in regulating this change in excitability after TENS [48]. On the peripheral side, the association of HDsEMG recordings with TMS enabled us to highlight the spatial distribution of MEPs, unveiling the change in corticospinal excitability within the pool of motor neurons innervating the FDI muscle [49, 50]. We observed consistent changes across electrodes together with similar coordinates of the centroids of MEP amplitudes before and after TENS (not reported here in the Results section), indicating that changes in corticospinal excitability were homogeneous within the pool of motor neurons.

We also tested how TENS impacts the excitability of spinal motor neurons through PIC. We used the delta F method that estimates the amplitude of PIC by calculating the hysteresis between the discharge rates of a ‘control’ motor unit at the recruitment and derecruitment times of a ‘test’ motor unit [34]. This method assumes that both ‘control’ and ‘test’ motor units receive a common synaptic input [34, 35]. Here, we showed that motor units from the FDI muscle receive a high amount of common synaptic input using the classic coefficient of determination between motor unit discharge rates (R^2^ = 0.93 ± 0.02 and 0.93 ± 0.03 before and after the session, respectively). We further confirmed this information with the significant level of coherence between motor units in a bandwidth from 0 to 5 Hz (**Figure 5**). It is also noteworthy that we used optimal timing differences between recruitment and derecruitment of ‘control’ and ‘test’ motor units to guarantee the full activation of PIC and limit the overestimation of delta F [25, 35]. We are therefore confident regarding the consistency of our delta F values to estimate PIC amplitudes. We found that the amplitude of PIC increased after a session of TENS (**Figure 3**). Numerous studies have found an acute increase in motor unit discharge rate during tendon vibration or TENS linked to the activation of PIC [10, 23, 51, 52]. Animal and human data showed that the repetition of these stimuli led to short-term plasticity referred to as ‘facilitation’ or ‘warm-up’ [10, 23, 51–53]. Molecular mechanisms behind this state involve voltage-gated calcium channels on the dendritic surfaces of spinal motor neurons mediated by the concentration of calcium ions [21]. However, ‘warm-up’ is a short-term state lasting several seconds. Here, we found an increase in PIC amplitude up to 5 min after the discontinuation of TENS, referred to as a ‘primed’ state. Recent work described an increase in motor neuron excitability at the spinal level with facilitation of the H-reflex after 20 min of TENS [13]. They proposed that the repeated stimulation of cutaneous afferent fibers may decrease Ia presynaptic inhibition, though they did not directly assess this parameter after the session of TENS. This could impact the amplitude of PIC as synaptic inhibition reduces PIC during isometric contraction [24, 54]. Alternatively, the increase in delta F values may also depend on the strength of monoaminergic input from the brainstem, which modulates PIC amplitude [55]. However, a previous study failed to demonstrate a change in brainstem activity following a session of TENS [48], and the current study was not designed to measure descending input from the brainstem. Therefore, future work should specifically investigate the potential sources of changes in PIC amplitude associated with a session of TENS.

We finally investigated how TENS alters the neural control of force by tracking the discharge properties of motor units across the session. We did not find a change in motor unit discharge properties, i.e., recruitment thresholds and discharge rate, or in the amplitude of common synaptic input to the FDI muscle (**Figures 4 and 5**). Moreover, force steadiness and the temporal variability of the cumulative spike train discharge rate remained unchanged after TENS. These results showed that a session of TENS is unlikely to induce short-term adaptations of force steadiness in healthy participants, whereas previous work found acute changes in force steadiness during TENS [12, 27, 56, 57]. An acute increase in the variability of common synaptic input likely caused the alterations of force steadiness observed elsewhere. Indeed, several studies have shown a positive correlation between the common modulation of motor unit discharge rates and the fluctuations of force exerted during an isometric contraction [1, 3, 5, 7]. Moreover, force steadiness was correlated to the variability of common synaptic input in healthy adults [5]. Similarly, we found in this study that motor units from the FDI muscle received a significant proportion of common synaptic input in the bandwidth of force fluctuation, with consistent levels before and after the session of TENS (**Figure 5**). Additionally, the temporal variability of the discharge rate of the cumulative spike train at low frequency did not change. Thus, TENS may acutely increase the variance in common synaptic input from sensory fibers [7, 27], an effect that disappeared once we discontinued the stimulation. Finally, it is noteworthy that the effect of TENS on force steadiness followed different trends than a previous study when comparing young and older adults[27]. In addition, TENS appeared to improve motor function in patients with multiple sclerosis in a clinical setting, while the authors did not find such improvements in healthy controls [41]. Therefore, we cannot extrapolate our results to other populations, and future experimental works remain necessary to understand how TENS may impact motor unit discharge dynamics and improve force steadiness in older adults or patients.

## 5. CONCLUSION

To conclude, TENS may only act as a priming method with healthy young participants. Here, ‘priming’ relates to stimuli that change neural excitability and, ultimately, impact subsequent motor tasks [13, 58]. We found in this study two potential sources of the primed modulatory effect of TENS on corticospinal excitability. First, the supplementary sensory input may reach supraspinal centers to alter the cortical representation of the FDI muscle and cortical excitability. Second, the gain between synaptic input and motor neuron output may change with an increase in PIC amplitude. It is noteworthy that these changes did not translate into an altered dynamic of motor unit activity during submaximal isometric tasks. Thus, the influence of afferent feedback on common synaptic input, indirectly estimated using the amplitude of motor unit coherence in the alpha band (5 – 15 Hz [59]), did not change after the session. Alternatively, the amplitude of the descending synaptic input may have changed. However, the non-linearity between motor neuron input and output due to neuromodulation prevents us from accurately estimating the level of synaptic input [21], and thus proving such explanation. Overall, increasing corticospinal excitability may have applications in a clinical setting. Indeed, several studies showed that priming techniques could enhance rehabilitation outcomes in patients with altered volitional control of movement (post stroke in [40]; spinal cord injury in [60]) or with sensory impairments (multiple sclerosis in [41]).

## Acknowledgments

We thank François Hug for his comments on the manuscript. We also thank SRALAB for funding this study.

## Data availability statement

The data that support the findings of this study are available from the corresponding author upon reasonable request.

